# At-level allodynia after mid-thoracic contusion in the rat

**DOI:** 10.1101/2020.08.10.240499

**Authors:** G. H. Blumenthal, B. Nandakumar, A. K. Schnider, M. R. Detloff, J. Ricard, J. R. Bethea, K. A. Moxon

## Abstract

The rat mid-thoracic contusion model has been used to study at-level tactile allodynia after spinal cord injury (SCI), one of the more common types of allodynia. An important advantage of this model is that not all animals develop allodynia and, therefore, it could be used to more clearly examine mechanisms that are strictly related to pain development separately from mechanisms related to the injury itself. However, how to separate those that develop allodynia from those that do not is unclear. The aims of the current study were to identify where allodynia and spasticity develop and use this information to identify metrics that separate animals that develop allodynia from those that do not in order study difference in their behavior. To accomplish these aims, a standardized grid was used to localize pain on the dorsal trunk and map it to thoracic dermatomes, providing for the development of a pain score that relied on supraspinal responses and separated subgroups of animals. Similar to human studies, the development of allodynia often occurred with the development of spasticity or hyperreflexia. Moreover, the time course and prevalence of pain phenotypes (at-, above-, or below level) produced by this model were similar to that observed in humans who have sustained an SCI. However, the amount of spared spinal matter in the injured cord did not explain the development of allodynia, as was previously reported. This approach can be used to study the mechanism underlying the development of allodynia separately from mechanisms related to the injury alone.

## Background

After a spinal cord injury (SCI), over 50% of individuals develop chronic neuropathic pain (CNP) (Burke et al., 2017) described as “severe or excruciating” in nearly half of all patients that experience it (Siddall et al., 2003). Unfortunately, CNP remains largely refractory to treatment and may be accompanied by comorbidities such as depression (Cairns et al., 1996), further reducing quality of life. A greater understanding of the mechanisms underlying CNP is essential to the development of more effective treatments (Cohen & Mao, 2014). However, before these underlying mechanisms can be explored, it is important that animal model used in these investigations include the relevant control group that undergoes a spinal cord injury but does not develop CNP. Moreover, these models should produce a similar prevalence and time course of pain phenotypes (no pain versus at-, above- or below-level) as observed in humans (Burke et al., 2012).

Several animal models exist to study CNP, and, mid-thoracic spinal cord contusion in the rat has been used to study central injury models due to its clinical relevance and ease at which it can be implemented (Metz et al., 2000; Sharif-Alhoseini et al., 2017). One of the most important advantages of this model is that not all animals develop allodynia and, therefore, this model could be used to study differences in underlying mechanism specifically related to pain separately from mechanisms related to the injury itself. Moreover, previous research found that mid-thoracic contusion in the rat results in the development of allodynia at multiple levels relative to the site of injury: above- (forepaws), at- (trunk), and below- (hindpaws) level allodynia (Hulsebosch et al., 2000; Lindsey et al., 2000), yet, differences between these subgroups of animals is less clear.

Thus, the primary aim of this study was to refine the assessment of at-level tactile allodynia in the rat model of mid-thoracic contusion in order to separate animals that develop allodynia from those that do not. The secondary aim was to describe the prevalence and onset of the different pain phenotypes (above-, at-, and below-level), to examine the time course of pain development and to determine if spared grey or white matter could be used to predict pain development. To accomplish these aims, a grid drawn on the dorsal trunk was mapped to thoracic dermatomes and the distribution of supraspinal responses to tactile stimulation of the trunk, forepaws and hindpaws was studied. Trunk allodynia was located just rostral to the level of the lesion and audible vocalizations and/or avoidance behaviors were the most informative to identify animals that develop allodynia. Trunk allodynia was the most prevalent pain phenotype, developing early, while fore- and hindpaw tactile allodynia was less common and developed later. Finally, spared matter within the cord did not correlate with behavioral measures of pain. Given similarities to human prevalence of CNP including distribution of pain phenotypes and time course of development as well as the availability of a control group of SCI animals that do not develop allodynia suggests this is a good model to study the mechanisms underlying the development of pain.

## Methods

### Subjects

One hundred and fifty-nine adult, female Sprague Dawley rats (225-250 g; Envigo) were used in this study. One hundred and thirty-eight rats received a moderate mid-thoracic spinal cord contusion, six received a laminectomy, and 15 non-injured animals were used to identify the location of the thoracic dermatomes. Of the contused animals, nine died during SCI surgery due to complications and three were removed from the study due to improper SCIs, defined as BBB scores greater than 20 at Week 1. Of the remaining contused animals, nine were used to assess locations on the trunk that, when stimulated, produced a painful response. These nine animals along with an additional 37 with a smaller grid were used to quantify the prevalence of evoked supraspinal responses to tactile stimuli on the trunk. A subset of these 46 animals, and 79 additional animals, were used to assess the impact of mid-thoracic spinal contusion on the development of trunk, forepaw and hindpaw allodynia (*N* = 117). Finally, to determine if the extent of damage in the cord was related to the development of pain, the amount of spared white matter was correlated to the number of supraspinal response in 11 of these animals.

All animals were maintained on a 12/12 hr light-dark cycle with food and water *ad libitum*. All experimental procedures were approved by the Drexel University and the University of California, Davis Institutional Animal Care and Use Committees (IACUC).

### Standardized Grid

To identify the location of thoracic dermatomes, and thereby localize trunk pain on the dorsal trunk relative to the location of the injury, a standardized grid was drawn on the dorsal trunk in a subset of animals while the animal was anesthetized with 2% isoflurane at least 24 hrs before testing. Each animal’s dorsum was shaved. To define the length of the trunk, the midpoint along a virtual line connecting the intertragic notches of the ears was connected to a point at the base of the tail and this distance was divided into 16 equally spaced grid rows. Next, four equally spaced grid columns were defined on each side of the vertebral column, from the midline to the lateral aspect of the dorsal trunk parallel to the knee for a total of eight columns (Fig. 1a). These columns and rows were draw on the animal’s skin to define the large trunk grid consisting of 128 grid squares, each approximately 1 cm^2^ due to the similar size of all animals. For additional behavioral testing of trunk allodynia in a larger group of animals, a smaller trunk grid with similar spacing localized to the region of the trunk most likely to develop pain (see Results) was used, which consisted of 40 grid squares, each approximately 1 cm^2^ (Fig. 1a).

**Figure 1:**
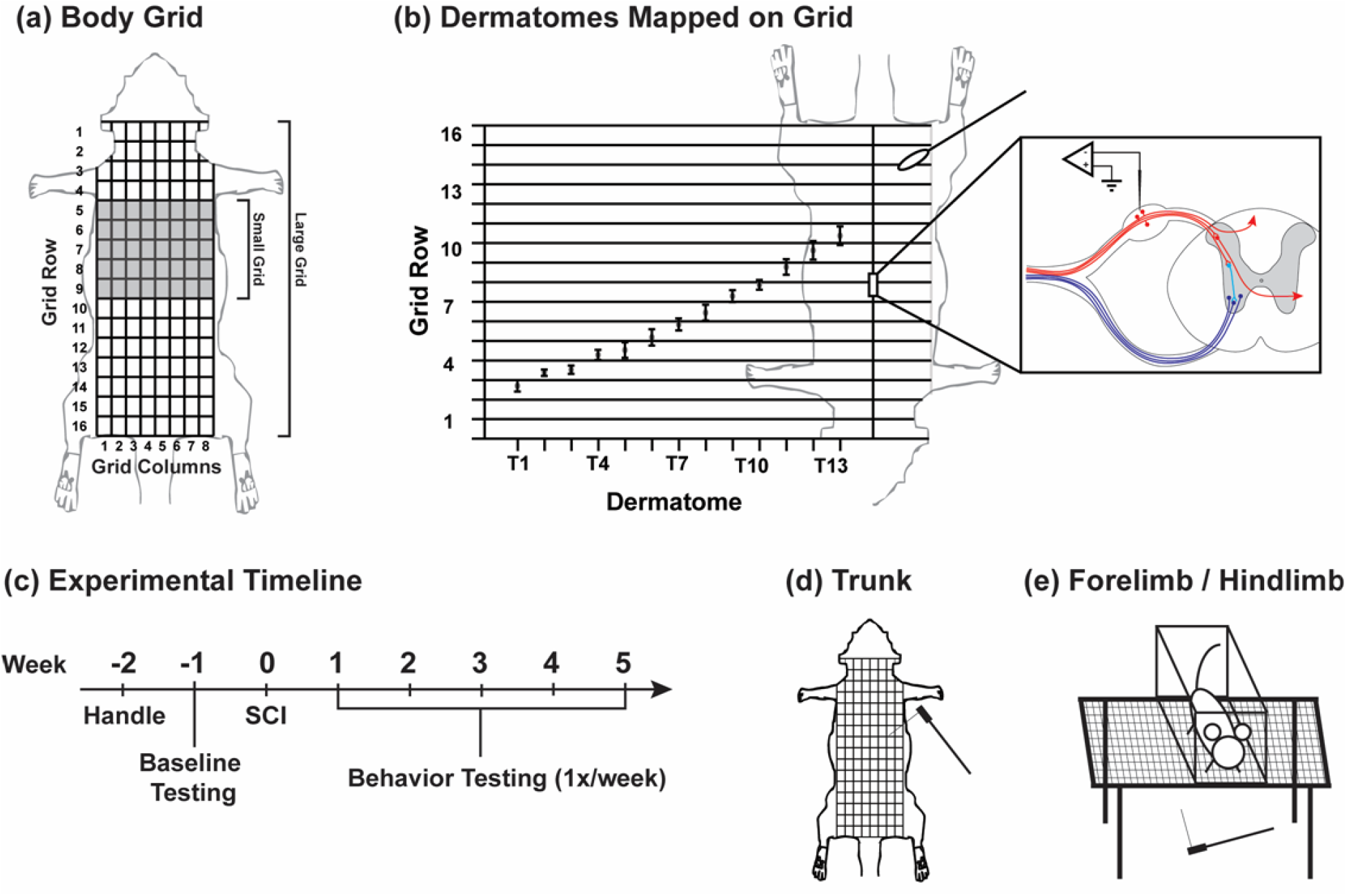
Experimental methods and timeline. (a) A large body grid was used to identify locations that when stimulated induce a painful response, while a small body grid was used to define at-level pain in a larger group of animals. (b) Grid locations were mapped to thoracic dermatomes by recording from multiple cells within each thoracic DRG and mapping their receptive fields relative to the grid. (c) Experimental timeline: Animals were handled and acclimated to each testing environment. Behavioral testing was performed once pre-SCI (baseline), and once a week for 5 weeks post-SCI. (d) A 26 g von Frey monofilament was used on the dorsum of the trunk to detect and quantify trunk tactile allodynia. (e) An electronic von Frey anesthesiometer was used to detect and quantify forepaw and hindpaw tactile allodynia.

### Thoracic Dermatome Map

To identify which dermatomes were likely to be associated with trunk allodynia after mid-thoracic SCI, thoracic dermatomes were identified in relation to the large trunk grid (Fig. 1b). Naïve uninjured animals were anesthetized with urethane (1.5 g/kg) via IP injection and maintained at a Stage III-3 anesthetic state (Friedberg et al., 1999). The skin and musculature overlying thoracic vertebrae T1-T13 were retracted carefully to avoid damage to any spinal nerves. The spinous processes, lamina, and transverse processes of selected vertebrae were removed unilaterally to expose the dorsal root ganglions (DRG). The spinal column was stabilized by attaching locking forceps to the transverse process immediately rostral and caudal to the selected vertebrae. A single high-impedance (4-10 MΩ) tungsten microelectrode (FHC Inc., Bowdoin, ME) was attached to a stereotaxic manipulator and positioned to a single DRG. A ground wire was placed in contact with the body cavity. The electrode was slowly inserted into the DRG as the neural signal was amplified (100X), band pass filtered (150 - 8000 Hz), digitized (40 kHz) (Plexon Inc., Dallas, TX), and monitored both on an oscilloscope and through audio speakers. Once a single unit was isolated, advancement of the electrode was paused, and light tactile stimulation was applied to the animal’s skin to determine the cell’s receptive field relative to the trunk grid (identifying which grid locations when given light tactile stimulation, resulting in the cell increasing its firing rate). The electrode was then advanced, and the process was repeated until another cell was identified or the electrode punctured through the DRG. Each DRG was sampled at least three times in different locations of the ganglion.

### Spinal Cord Contusion

Animals were anesthetized with ketamine (63 mg/kg), xylazine (5 mg/kg) and acepromazine (0.05 mg/kg) via IP injection. Animals were considered sufficiently anesthetized with the absence of a toe pinch reflex. The skin and musculature overlying the spinal column was retracted from spinal levels T4 to T12 and a laminectomy was performed at vertebral level T10. The spinal cord was stabilized by securing locking forceps to the transverse processes of T9 and T11. SCI rats received a moderate contusion injury at vertebral level T10 using the Infinite Horizon impactor device (Precision Systems and Instrumentation, LLC; Fairfax Station, VA) with 150 kdynes of force and a 1 s dwell time. The musculature was then sutured in layers and the skin was closed with wound clips. Laminectomy controls underwent the same procedure, except the spinal cord was not impacted. Animals were post-operatively hydrated with saline (7 ml), prophylactically administered an antibiotic (enrofloxacin, 5 mg/kg), and allowed to recover on a heated water pad. Animals were administered fluids and antibiotics once daily for seven days and bladders were manually expressed twice daily until they regained autonomic bladder control.

### Behavioral Testing

Prior to locomotor assessment and allodynia testing, animals were handled and habituated to the behavioral testing environments. This consisted of placing each animal in the open field (locomotor testing environment; 15 mins) and the von Frey testing cage (forepaw and hindpaw allodynia testing environment; 30 mins), as well as cradling each animal in the testers’ forearm (trunk allodynia testing environment; 30 mins), once a day for three days. After habituation, behavioral testing was conducted on all animals pre-operatively to establish baseline measures, and then once a week post-operatively for five weeks (Fig. 1c).

Locomotor functional recovery was measured using the Basso, Beattie, and Bresnahan (BBB) locomotor rating scale (Basso et al., 1995). Animals were placed in an open field (76.20 × 91.44 cm) and were observed by two trained experimenters blinded to the animals’ experimental condition for 4 mins. Each hindlimb was assessed for the presence of joint movements, weight support, quality of stepping, forelimb-hindlimb coordination, paw placement, and toe clearance and these observations were converted into a BBB score for each hindlimb. Scores on this scale range from 0 to 21, where a score of 0 represents a complete paralysis of the hindlimbs, while a score of 21 represents the locomotor function of an uninjured rat.

One day prior to the trunk allodynia test, animals were briefly placed under isoflurane anesthesia, the dorsal trunk was shaved, and the trunk grid was drawn on the skin. During the trunk testing session, the experimenter draped an absorbent pad across their forearm and placed the animal on the pad, unrestricted such that it was free to walk back and forth across the experimenter’s forearm. The animal supported its own weight on all four limbs for the entirety of the testing session. A 26 g force von Frey filament (Stoelting; Wood Dale, IL) was applied perpendicularly to the dorsal surface of the trunk until the filament bent (Fig. 1d). The filament was randomly applied to the center of each trunk grid square (large grid - 128 squares, smaller grid - 40 squares) until the entire grid was stimulated, with an interstimulus interval of at least 10 s. This process was repeated a total of five times. A 26 g force was selected because it has been documented to be a normally non-noxious tactile stimulus for similarly sized animals (Hulsebosch et al., 2000). Observable aversive supraspinal responses were indicative of pain and included audible vocalizations, biting at the filament, licking the point of stimulation, looking at the filament, and avoidance behavior in direct response to the stimulus. The stimulated trunk grid location that elicited the response and the type of response was documented. Animals generally never evoked more than one type of response upon a single stimulation. In addition to evoking supraspinal responses, stimulation of the trunk also evoked hindlimb movements without supraspinal responses that were unrelated to voluntary movements (Baastrup et al., 2010).These response were considered spastic and the stimulus location that elicited them was noted.

For forepaw and hindpaw allodynia testing, standard methods were used (Ängeby Möller et al., 1998). Briefly, animals were placed in a Plexiglas chamber (10.16 × 25.40 × 10.16 cm cage) with a wire mesh bottom and were allowed to acclimate to the environment for at least 20 mins before testing began. An acclimated animal displayed little to no movement or exploratory behavior. For each paw, an electronic von Frey filament (Ugo Basile, Gemonio, Italy) with a stiff metal tip was slowly applied to the plantar surface of the paw between the paw pads at a constant rate as the device measured the force which was being applied. The force at which the animal quickly withdrew its paw was recorded as well as any supraspinal responses made by the animal during the trial (Fig. 1e). Five stimulations were applied to each paw per session with an interstimulus interval of at least 30 s. To ensure accurate withdrawal thresholds, testing was only carried out if animals had the ability to bear weight on all limbs.

### Histology

Histological verification of the spinal cord lesion was conducted on subset of animals. At the conclusion of behavioral testing five weeks post-SCI, animals were transcardially perfused with cold saline followed by 4% paraformaldehyde (pH 7.4). During spinal cord tissue removal, the vertebral level of the lesion site was confirmed. Tissue was post-fixed in 4% paraformaldehyde for 24 hrs and placed in 30% sucrose until the tissue sank to the bottom of the specimen container, indicating that the tissue had been cryoprotected. A 14 mm section of spinal cord surrounding the lesion site was dissected and frozen in Shandon M1 embedding matrix (Thermo Fisher Scientific, Waltham, MA). 25 μm coronal sections of cord were collected using a freezing microtome and every 20^th^ slice was mounted onto charged slides (Thermo Fisher Scientific premium frosted microscope slides) to preserve 500 μm spacing between section. Sections were air dried overnight. To stain, the slides were dehydrated in increasing concentrations of ethanol baths (75%, 95%, 100%) for 3-6 mins each, cleared using Citrisolv (DeconLabs Inc., King of Prussia, PA) for 20 mins, rehydrated in decreasing concentrations of ethanol baths (100%, 95%, 75%) for 3-6 mins each, and were then rinsed with distilled water. The slides were stained for myelin using a Cyanine R / FeCl_3_ solution for 10 mins. Slides were rinsed and placed in differentiation solution for 1 min using 1% aqueous NH_4_OH. After additional rinsing, slides were stained for Nissl in a Cresyl Violet solution for 20 mins, rinsed, and dehydrated once again using ethanol. Slides were coverslipped using Vectashield mounting medium (Vector Laboratories, Burlingame, CA) and digital images were taken of each section 24 hrs later.

### Data Analysis

#### Dermatome map analysis

For each animal, a thoracic dermatome was identified as the union of all trunk grid locations that, when stimulated, modulated the firing rate of any cell recorded from a single DRG. Dermatome width, defined as the rostro-caudal extent of each dermatome measured in trunk rows, was calculated for each DRG sampled. Additionally, the central position of each dermatome was calculated, defined as the point on the trunk grid at the center of each dermatome’s width. Dermatome widths and central positions were then averaged across all animals, and averaged thoracic dermatomes were defined by taking the average dermatome width centered on the average dermatome center position.

#### Behavioral assessment

The frequency, type, and location of supraspinal responses to tactile stimulation of the trunk grid were noted and used to refine trunk pain assessment. To assess the rostro-caudal extent of allodynia, the number of supraspinal responses across each grid row for each animal was tallied, separately for each week. The values at Week 5 were used to define a pain score for trunk tactile allodynia (see Results). Similarly, to assess development of spasticity the sum of spastic responses across each grid row was calculated at Week 5.

The development of forepaw and hindpaw allodynia was evaluated at each week post SCI. Similar to previous studies, the median force of the five trials performed on each paw during a testing session was considered to be the withdrawal threshold for that paw in that testing session. There was not a significant difference in baseline withdrawal threshold between the left and right forepaws [*t*(47) = 1.42, *p* = .16], or hindpaws [*t*(45) = .13, *p* = .89], so paw withdrawal thresholds were averaged between the left and right paws. Withdraw thresholds were then normalized to the animal’s baseline score. An animal was considered to have tactile allodynia in the forepaws or hindpaws if the withdrawal threshold was reduced by at least 50% compared to their baseline withdrawal threshold, and the animal exhibited a supraspinal response during stimulation (Detloff et al., 2013). If an animal had a >= 50% decrease in withdrawal threshold at Week 5 compared to baseline, but did not exhibit supraspinal responses during stimulation, it was considered to have hyperreflexia, but not allodynia Finally, to assess locomotor recovery, the BBB scores from the left and right hindlimb of each animal were averaged together such that each animal had a single BBB score for each week.

#### Histological analysis

To determine if there was an association between spared matter around the lesion site and the presence of allodynia, the amount of spared white and grey matter in each section was calculated using Image J Software (NIH, Bethesda, MD) (Schneider et al., 2012). Tissue was considered spared if staining was uniform and it was absent of extensive cellular debris or vacuoles. All measured sections were then normalized to the section with the largest amount of total spared tissue and converted to a percentage of spared tissue. The section with the least amount of total spared tissue was considered the lesion epicenter.

Because hemispheric asymmetries in the ventrolateral funiculus (VLF) have been suggested to occur more often in animals that develop pain compared to those that do not (Hall et al., 2010), the relationship between asymmetries in the amount of spared grey matter and the number of supraspinal responses was assessed using a similar approach. Briefly, the cord was divided into quadrants by drawing a vertical line and a horizontal line through the central canal. To isolate the VLF from the ventromedial funiculus, the lower quadrants were then further divided by a line drawn from the tip of each ventral horn to the edge of the section (Fig. 6a). As in Hall et al., 2010, if the ventral horns were damaged to such an extent that they could not be identified, their medial border was estimated by drawing a line from the central canal to the ventral edge of the section at a 30° angle, which is approximately the angle of the line drawn on a naïve cord (Fig. 6b).

### Statistical Analysis

Analysis of supraspinal responses to trunk tactile stimulation at Week 5 was used to refine our model of trunk allodynia. The distribution of the number of supraspinal and spastic responses per row of the large grid was used to assess the location of trunk allodynia and define a smaller grid. The distribution of the different types of supraspinal responses across the smaller grid was used to further develop a method to separate animals with allodynia from those that did not develop allodynia (see Results). Chi-square test were used to assess the importance of supraspinal responses to distinguish allodynia from hyperreflexia in response to paw stimulation.

Using our operational definition of trunk allodynia, differences in behavioral measures between animals that developed allodynia compared to those that did not were compared over time using a repeated measures restricted maximum likelihood estimation linear mixed model with a Greenhouse-Geisser correction. Where appropriate, post-hoc Sidak pairwise comparisons were performed at a significance level of 0.05. This procedure was implemented to prevent list wise deletion due to missing data. All statistical analyses were conducted using GraphPad Prism 8.0.2 for Windows (GraphPad Software, San Diego, CA).

To assess the effect of the amount of spared white and grey matter on the development of trunk allodynia, the percentages of spared white and grey matter were separately averaged across animals for the sections located at the same distance from the lesion epicenter. Spared tissue across the lesion site in animals with trunk allodynia were compared to that of animals with no allodynia anywhere. The percent of total spared white or grey matter along the entire lesion site between groups was compared using the same repeated measures statistical test as that used for differences in behavioral measures. Finally, to evaluate if asymmetry of spared white or grey matter in the VLF near the lesion epicenter could account for the development of pain, the percent difference of spared matter between the VLFs of the right and left side of the cord were compared and correlated to number of supraspinal responses per animal using Pearson correlation tests.

## Results

### Localization of Trunk Allodynia and Spastic Responses

The full trunk grid was used to localize, and subsequently quantify, responses evoked by innocuous tactile stimulation. Of the 10 animals that were tested for trunk allodynia using the full trunk grid, seven animals exhibited supraspinal responses elicited primarily from stimulation of rostro-caudal grid rows 5-9. Electrophysiology determined that these grid rows correspond to spinal dermatome levels T4-T10 and include part of dermatome T11 (Fig. 2a). In fact, all animals that exhibited supraspinal responses responded to stimuli within the T4-T11 dermatomes. A subset of these animals had larger areas that elicited pain responses, predominately avoids, expanding into upper thoracic and cervical dermatomes. Very few supraspinal responses were elicited below the level of the lesion. Therefore, a moderate T10 spinal cord contusion consistently produced painful responses at, and immediately rostral to, the site of the lesion, with the majority of the responses defining a painful region up to six dermatomal levels above the lesion site.

**Figure 2:**
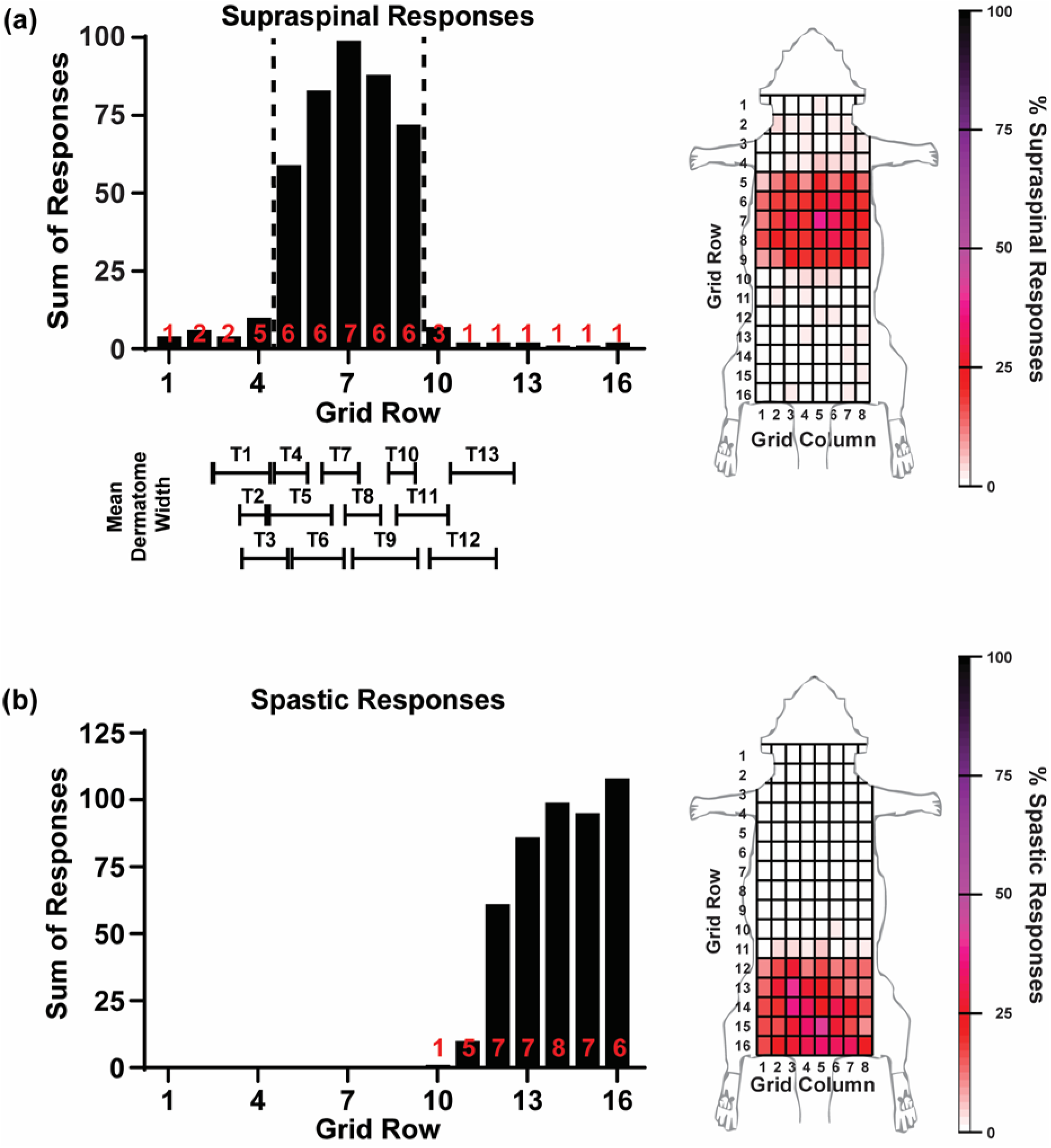
Localization of pain and spasticity. (a) Distribution of supraspinal responses to tactile stimulation of the dorsal trunk of the rat by grid row. The number of animals from which supraspinal responses were detected in each grid row are indicated in red on the histogram. A mapping of the grid rows to the thoracic dermatomes is shown below. Note that there is considerable overlap between adjacent dermatomes. (b) Heat maps displaying the locations and percent of possible evoked responses for each supraspinal response. In each grid square, 100% = 50 responses or five stimulations per square x 10 animals). (c) Distribution of spastic responses to tactile stimulation of the dorsum of the rat. (d) Heat map displaying the locations and percent of possible evoked spastic responses.

Spastic or rapid extension of the hindlimbs in response to trunk stimulation was also observed. Tactile stimulation to trunk grid rows 11-16 elicited these spastic responses, which correspond to dermatome levels T12 and below, extending into lumbar dermatomes (Fig. 2b). Responses were mainly bilateral and were in response to stimulation across the mediolateral extent of the trunk dorsum. These spastic responses were not accompanied by supraspinal responses. While locations that produced spastic responses were rarely colocalized (occurring in only one animal), spastic and supraspinal responses did occur in the same animal: of the 10 animals tested with the full grid, two had supraspinal responses only, three had spastic responses only, and five had both supraspinal and spastic responses. Because the majority of supraspinal responses occurred mainly rostral to the lesion site (T4-T11) and were generally not co-localized with spastic responses (below T11), it was determined that a smaller grid could be used to identify painful responses.

This smaller grid was used on a larger sample of animals to determine the best behavioral markers of trunk allodynia (Table 1). As expected, sham SCI animals elicited few supraspinal responses to non-noxious tactile stimulation. At Week 5, the most common supraspinal response in SCI animals was vocalization, comprising 52.71% of all responses, followed by avoid, which comprised 28.93% of responses. To localize these responses, the stimulus location was mapped to the grid (Fig. 3a). Vocalizations were located across the entire grid, which encompassed dermatomes T4-T11, while avoids were concentrated more medial and anterior. Because lick, look, and bite responses occurred relatively infrequently (9.11%, 7.79% and 1.45%, respectively), vocalization and avoid responses were sufficient to discriminate animals that developed pain from those that did. Therefore, the percentage of vocalization and avoids for each animal was used as a pain score (200 stimuli per animal).

**Table 1:** Contused animals primarily vocalize and elicit avoidance behavior in response to tactile stimulation of the trunk. The incidence of their evoked supraspinal responses increases over time. In contrast, laminectomy animals elicited few supraspinal responses and the incidence of their responses did not change over time.

| Response | Baseline |  | Week 1 |  | Week 2 |  | Week 3 |  | Week 4 |  | Week 5 |  | % |
| --- | --- | --- | --- | --- | --- | --- | --- | --- | --- | --- | --- | --- | --- |
|  | Sham | SCI | Sham | SCI | Sham | SCI | Sham | SCI | Sham | SCI | Sham | SCI |  |
| Vocalize | 2 | 4 | 2 | 55 | 0 | 21 | 4 | 164 | 0 | 137 | 2 | 399 | 52.7 |
| Avoid | 0 | 9 | 1 | 62 | 0 | 47 | 0 | 149 | 0 | 204 | 0 | 219 | 28.9 |
| Look | 0 | 19 | 3 | 36 | 0 | 30 | 7 | 39 | 2 | 27 | 0 | 59 | 7.8 |
| Lick | 0 | 6 | 0 | 0 | 0 | 2 | 0 | 49 | 0 | 82 | 0 | 69 | 9.1 |
| Bite | 0 | 3 | 0 | 4 | 0 | 0 | 0 | 0 | 0 | 3 | 0 | 11 | 1.5 |

*Note.* Sham = laminectomy animals (n = 6) and SCI = contused animals (n = 47). % indicates the percentage of all supraspinal responses that is comprised of by that type of response, at Week 5, in contused animals.

**Figure 3:**
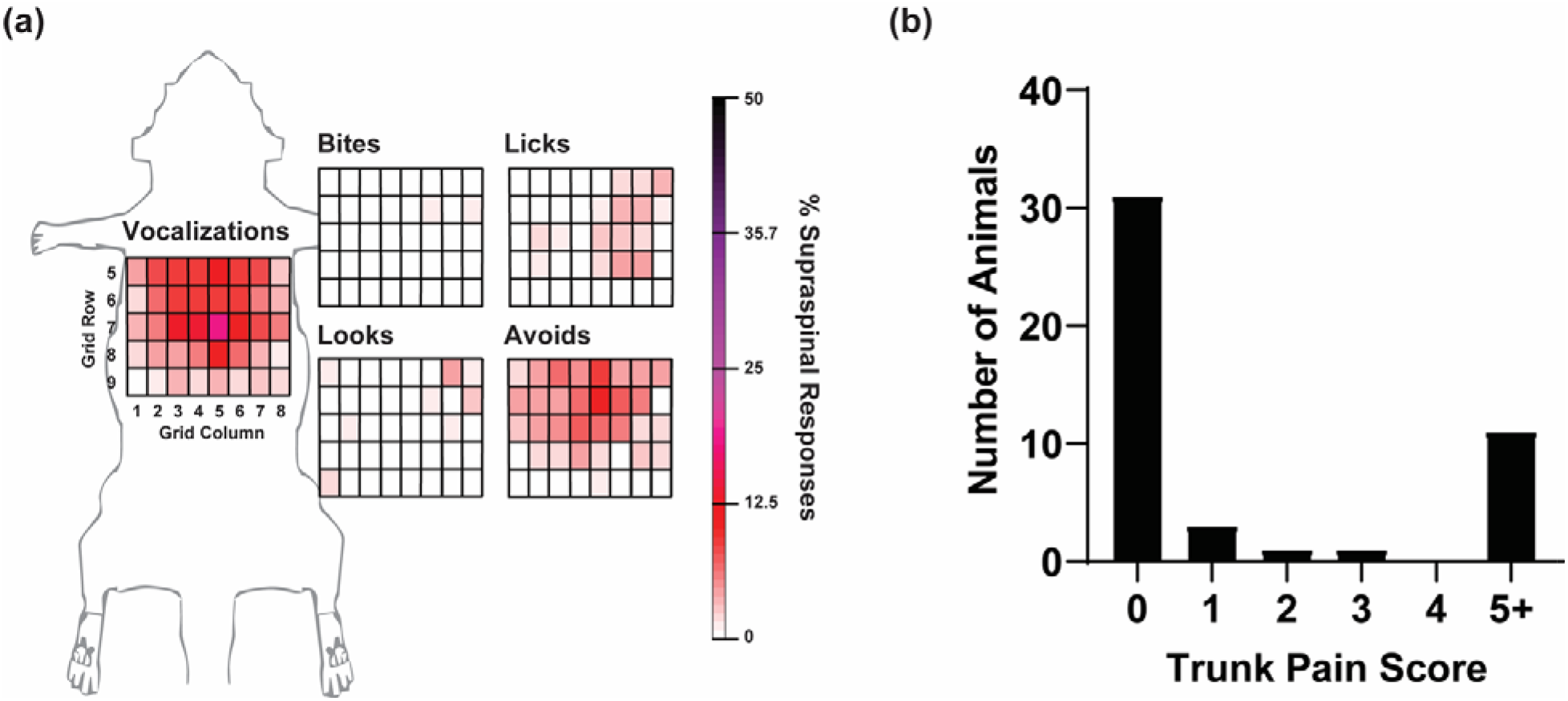
Characterization of at-level pain. (a) Heat map displaying percentage of each type of supraspinal response for each location and type of supraspinal responses evoked from stimulation of the smaller grid. (b) Distribution of pain scores. The pain score is defined as the sum of vocalization and acoid responses divided by the total number of stimulations (200). From this distribution, a score >=5 was used to distinguished animals with at-level allodynia from animals without at-level allodynia.

To identify a threshold that could separate animals that develop tactile allodynia from those that do not, the distribution of pain scores in the smaller grid across all animals was evaluated (Fig. 3b). The majority of animals had a pain score of zero, five animals had a pain score less than or equal to three, and 11 animals had a pain score greater than or equal to five (μ=26.95 +/− 17.71%). The average raw number of vocalize + avoid responses per animal was 53.91 +/− 35.41% occurring in an average of 20.91+/−7.80 grid locations. For those animals with a pain score of at least five, only two had a pain score less than 10 and over half had a pain score greater than 25 (Fig. 3b). Because animals that do not develop tactile allodynia are highly unlikely to produce a supraspinal response to a non-noxious stimulus, a threshold of five for the pain score can differentiate spinal injured animals that develop allodynia from those that do not.

To test if restricting the pain score to only include vocalization and avoid responses misrepresented the likelihood of identifying cases of allodynia, we compared our threshold pain score using only vocalizations and avoids to a threshold using all supraspinal responses (i.e., vocalize, bite, lick, look, avoid). We found that the same 11 animals would have been considered to have allodynia, therefore, not using bites, licks or looks did not affect whether an animal was considered to develop pain. Moreover, the percentage of all other evoked supraspinal responses (i.e. look, ick, bite) for animals with trunk pain was low (μ=5.72+/− 6.61), suggesting that these supraspinal responses may not be the best indicator of pain, and that using only vocalizations and avoids is sufficient to asses tactile allodynia in the trunk. Therefore, a pain score of at least five, calculated using vocalizations and avoids at Week 5, was used to distinguish animals that developed trunk allodynia from those that did not.

The same set of contused animals were also tested for the presence of spasticity of the hindlimbs in response to trunk stimulation at Week 5. In addition to testing within the small grid, animals were stimulated a total of 20 times on each side of the dorsal trunk between the caudal border of the small grid and the tail in random locations. Almost half of the animals (21 of 46) had at least one spastic response. Of the 541 total spastic responses detected, only five responses from two animals were elicited from within the small grid, whereas all other responses were elicited from between the caudal border of the small grid and the tail. Over half of the animals with allodynia (6 of 11) also had spastic responses, whereas only 28.57% (6 of 21) of animals with spastic responses also had trunk allodynia. Supraspinal and spastic responses were never elicited from the same trunk location in the same animal. Therefore, the trunk locations which elicit spasticity of the hindlimbs upon stimulation arise caudal to the region of trunk pain development, and the development of one does not require nor exclude development of the other.

### Animals Unlikely to Have Tactile Allodynia at Multiple Levels

To better understand the development of tactile allodynia in this model, a larger group of animals was tested for trunk allodynia using our pain score threshold developed above as well as forepaw and hindpaw allodynia. Each animal was classified into one of eight pain phenotype groups depending on the levels where allodynia developed (Fig. 4). An animal was more likely to develop allodynia at only one level (27.35% of animals: 18.80% trunk alone, 5.13% forepaw alone, 3.42% hindpaw alone) than at multiple levels (6.83% of animals: 2.56% forepaw + hindpaw allodynia, 1.71% trunk + forepaw allodynia, 2.56% trunk + hindpaw, 0% trunk + forepaw + hindpaw allodynia). Over half of animals (65.90%) did not display any supraspinal responses to forepaw, hindpaw or trunk tactile stimulation, suggesting that they did not develop tactile allodynia. In summary, 23.07% of animals developed trunk, 9.40% developed forepaw, and 8.54% developed hindpaw allodynia with 34.18% developing allodynia in at least one region.

**Figure 4:**
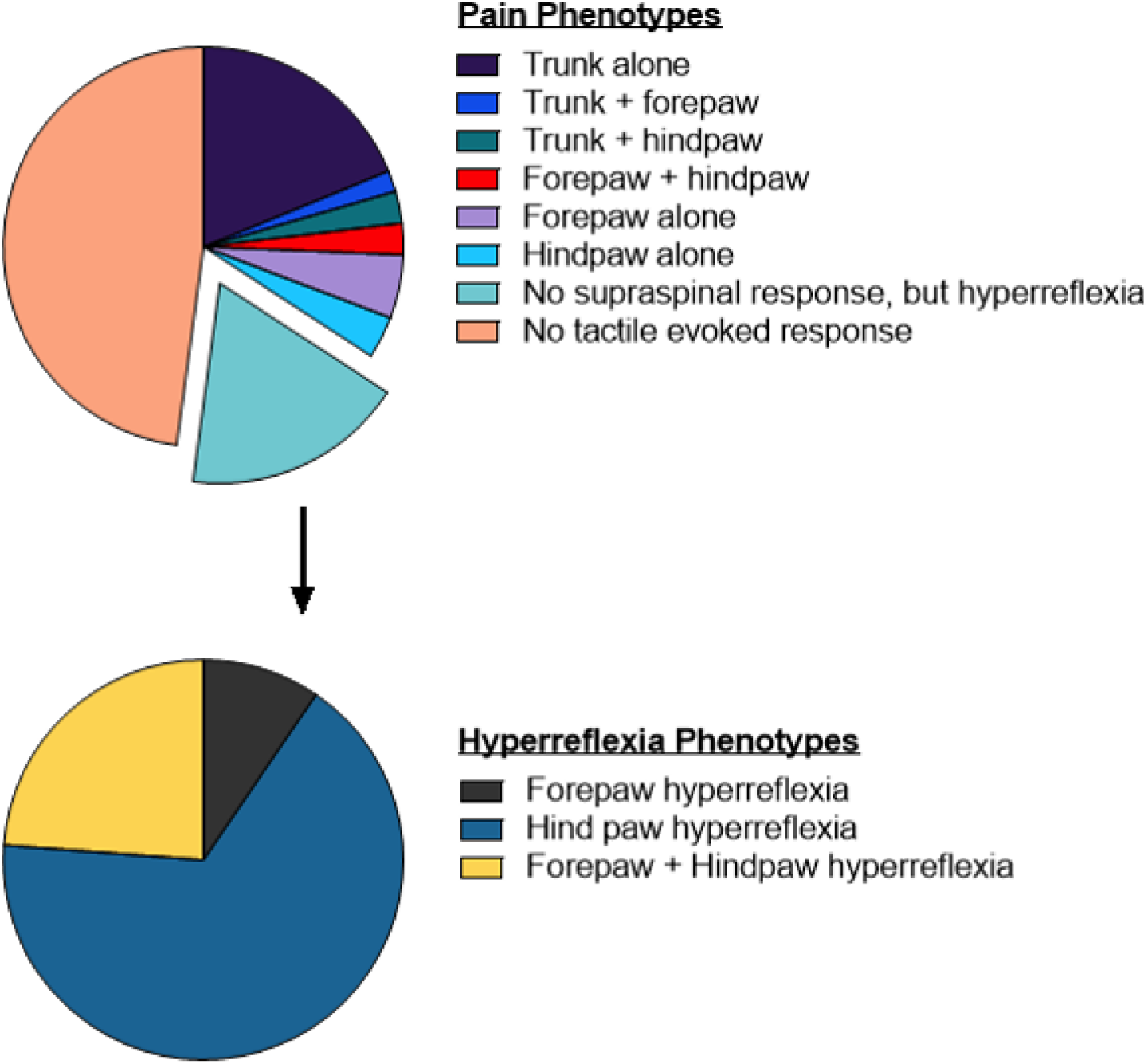
Frequency of Pain Phenotypes. Pain phenotypes were determined for each animal at Week 5 and were assigned based on whether the animal had at-, above-, below-level allodynia, or some combination of the three. For animals that had no supraspinal responses but demonstrated hyperreflexia in the limbs, the frequency of hyperreflexia for forepaw, hindpaw, or both are broken out in the second pie chart.

Of the animals that did not display supraspinal responses during forepaw or hindpaw tactile stimulation, 17.95% displayed hyperreflexia (i.e., a 50% or greater reduction in forepaw and/or hindpaw withdrawal threshold). In fact, the overall proportion of animals that developed hyperreflexia in the hindpaws (13.59%) was greater than the proportion of animals that developed hindpaw allodynia (3.54%) (*X*^2^ (1, *n* = 117) = 6.02, *p* = .01) with no difference between the overall number of animals that developed forepaw hyperreflexia (9.34%) and the number that developed forepaw allodynia (5.41%) (*X*^2^(1, *n* = 117) = 1.07, *p* = .30).

### Trunk Allodynia Develops Early Post Injury

To assess the effect of developing allodynia on locomotor recovery, the BBB scores of animals determined to have allodynia at any level (forepaw, trunk, and/or hindpaw) were compared to animals that did not develop allodynia anywhere. As expected, BBB scores decreased immediately after SCI, but then increased with each successive week post-SCI for both groups (main effect of week: *F*(2.61, 194.70) = 239.4, *p* < .001; main effect of group: *F*(1, 76) = 0.44, *p* = .88; interaction *F*(5, 373) = 1.06, *p* = .46, Fig. 5a). This suggests that there is not a relationship between locomotor recovery and pain development.

**Figure 5:**
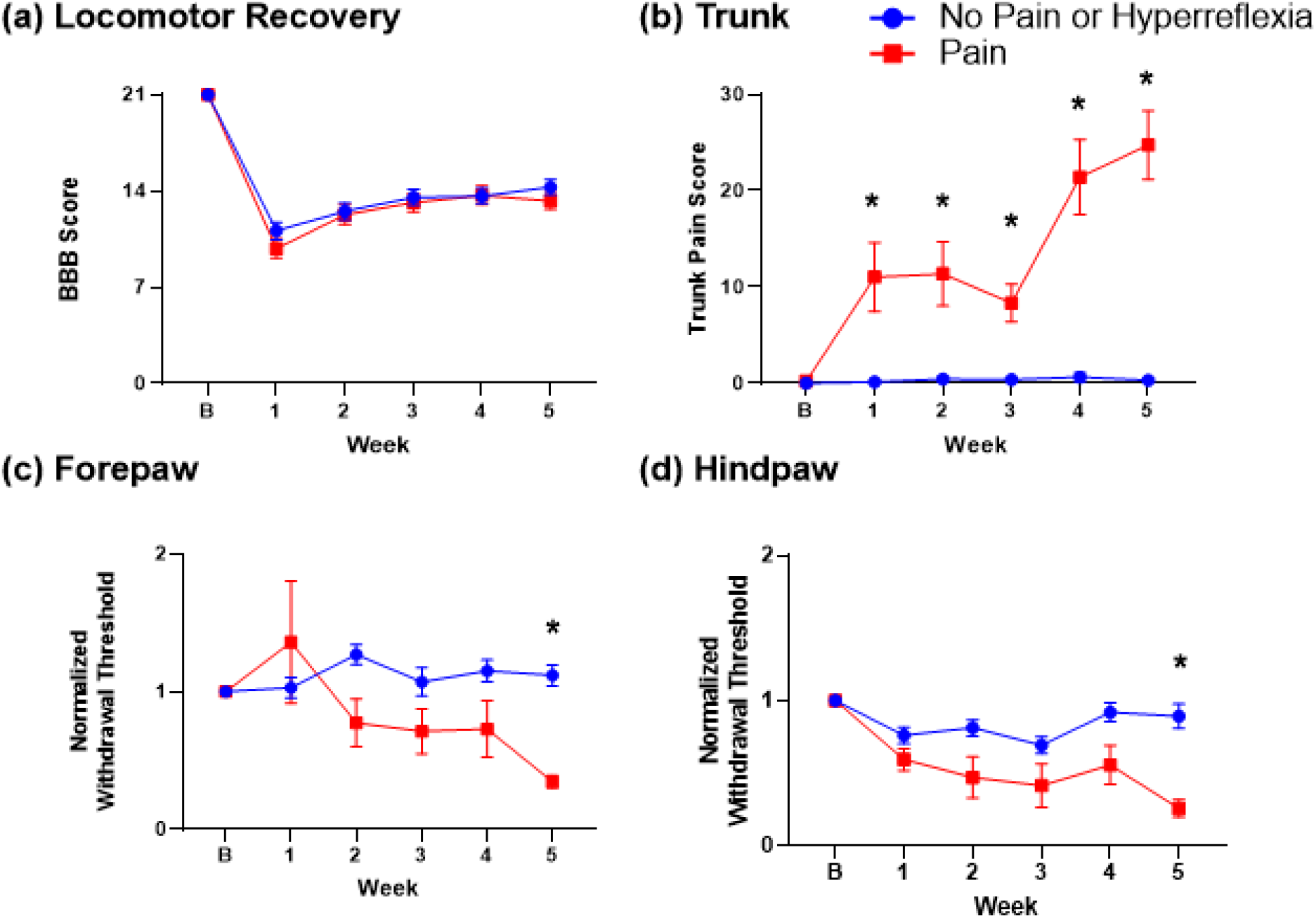
Temporal development of allodynia. Changes in behavior were compared across weeks for animals the developed allodynia and those did not. (a) Locomotor ability (BBB score) between animals that developed allodynia at any level (*n* = 40) was compared to that of animals that did not develop allodynia anywhere (*n* = 38). (b) The percentage of vocalizations was compared between animals that developed at-level allodynia (*n* = 29) and the animals that did not develop allodynia anywhere. (c) & (d) Withdrawal thresholds for forepaws and hindpaws of animals that developed forepaw (*n* = 10) or hindpaw (*n* = 8) allodynia, respectively, were compared to the animals that did not develop allodynia anywhere. For all plots (a-d) the no pain group was the same group of animals that did not develop allodynia anywhere (*n* = 38). Animals were classified into pain phenotypes at Week 5. B = baseline before SCI, numbers indicate weeks post SCI. \**p*<0.05.

To assess development of the different phenotypes of allodynia over time, the trunk pain score, and the hindpaw and forepaw withdrawal thresholds were compared separately across weeks between animals that developed allodynia at each level and those that did not develop allodynia at any level. The trunk pain score for animals that developed trunk allodynia significantly increased over time but not the score for animals that did not develop allodynia [main effect of week: *F*(3.39, 213.90) = 17.24, *p* < .001; main effect of group: *F*(1, 65) = 48.70, *p* < .001; interaction: *F*(5, 315) = 16.09, *p* < .001]. Importantly, the pain score significantly differed between groups starting in Week 1 and continued through Week 5 with the difference becoming greater with each successive week (Fig. 5b). Similarly for forepaw allodynia (*n* = 10), withdrawal thresholds for animals that developed allodynia decreased overtime while withdrawal thresholds for animals that did not develop allodynia remained stable [main effect of week: *F*(3.76, 166.90) = 4.05, *p* < .01; main effect of group: *F*(1, 46) = 6.50, *p* < .05; interaction [*F*(5, 222) = 6.33, *p* < .001]. However, differences between groups were apparent only at Weeks 2 and 5 (Fig. 5c). For hindpaw allodynia (*n* = 8), there were again significant differences between groups [main effect of week: *F*(3.66, 156.60) = 7.77, *p* < .001; main effect of group: *F*(1, 44) = 11.58, *p* < .005, interaction: *F*(5, 214) = 3.34, *p* < .01]. Withdrawal thresholds for animals that develop hindpaw allodynia decreased with time becoming significantly different from those that did not develop allodynia at Weeks 4 and 5 (Fig. 5d). Taken together these data suggest that trunk allodynia emerges early, while forepaw and hindpaw allodynia emerge at later time points after SCI. This is consistent with the development of chronic neuropathic pain in human SCI patients where hindpaw pain was found to have a later onset than trunk pain (Finnerup et al., 2014).

### Trunk Allodynia is Not Related to Differences in Lesion

To understand if differences in spared tissue (left versus right) were associated with trunk allodynia, the amount of spared grey and white matter across the lesion site was compared in a subset of animals with trunk allodynia (*n* = 7) and a subset of animals with no allodynia anywhere (*n* = 5). While thoracic sections of spinal cord distal to the lesion site (~7 mm) appeared completely undamaged (Fig. 6a), sections from the lesion epicenter generally suffered extensive damage, with a complete absence of grey matter and a small amount of spared white matter usually found along the ventral periphery of the cord (Fig. 6a). However, along the entire lesion site, there were no differences between the percent of total spared white or grey matter between pain groups [white matter main effect of group: *F*(1, 9) = 0.50, *p* = .50; main effect of distance from the epicenter: *F*(27, 186) = 24.94, *p* < .0001; interaction: *F*(27, 186) = 0.73, *p* = .83, Fig. 6c; grey matter main effect of group: *F*(1, 9) = 1.291, *p* = .96; main effect of distance from the epicenter: *F*(2.76, 17.71) = 29.32, *p* < .001; interaction: *F*(28, 180) = 0.58, *p* = .96, Fig. 6d]. This suggests that total sparing is unlikely to predict the development of allodynia. Moreover, we found no correlation between trunk pain score and spared white matter in the ventrolateral funiculus [*r(*9) = .06, *p* = .89 at the epicenter; or 1 mm rostral *r*(9) = .03, *p* = .92] (Fig. 6e), nor any correlation between trunk pain score and asymmetry of the funiculi at the lesion epicenter [*r(*9) = -.25, *p* = .46, or 1 mm rostral *r*(9) = .17*, p* = .61] (Fig. 6f). These results suggest that the amount of spared tissue at the lesion site does not explain development of trunk allodynia.

**Figure 6:**
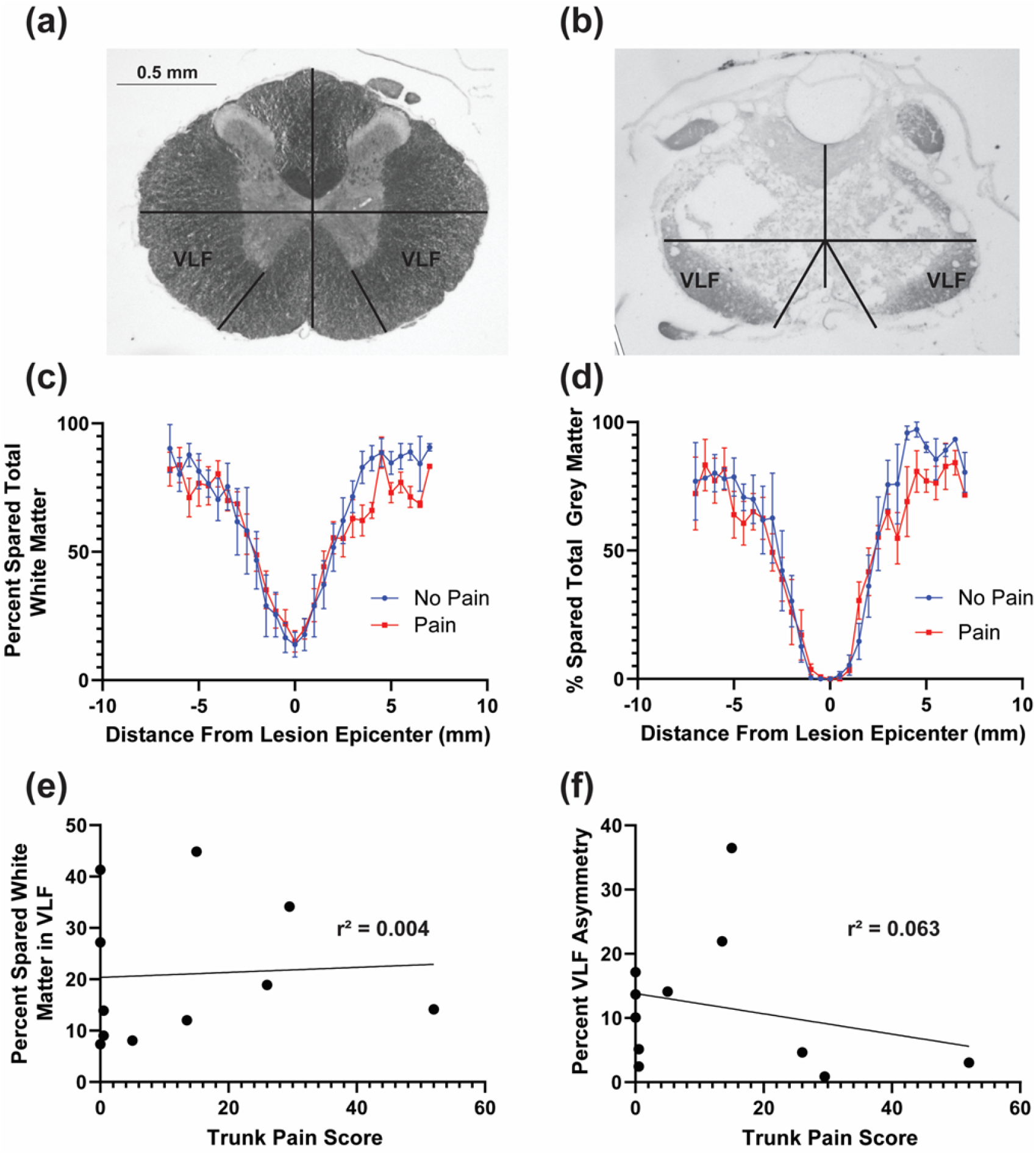
Assessment of spared white and grey matter. (a) An example of an undamaged histological section distal from the lesion site noting the area estimated to be the ventrolateral funiculi. (b) If the section had little spared matter, the locations of the ventrolateral funiculi were estimated (see Methods). (c) & (d) The amount of spared white or grey matter across the lesion site of animals that developed trunk allodynia was compared to animals that did not develop allodynia and no differences were found. (e) & (f) No correlation was found between the amount of spared white matter in the ventrolateral funiculus (VLF) or the degree of asymmetrical sparing of the VLF at the lesion epicenter (data not shown) or 1 mm rostral to the lesion and the trunk pain score.

## Conclusions

The mid-thoracic spinal contusion model of SCI is an important model for the study of CNP. To provide more insight into the impact of this injury on the development of allodynia, the assessment of trunk, or at-level, allodynia was refined. Tactile allodynia was found to occur at and immediately rostral to the lesion, and in distinct dermatomes from spastic responses due to trunk stimulation. Furthermore, the presence of allodynia could be discriminated by assigning animals a pain score based on the percentage of vocalization and avoid responses elicited, without consideration of other supraspinal responses. Of the animals that developed allodynia, the majority developed at-level (trunk) allodynia and this at-level allodynia developed earlier than below-level (hindpaw) or above level (forepaw) allodynia. These findings are similar to the prevalence and time course of pain phenotypes (no pain versus at-, above- or below-level) observed in humans (Burke et al., 2012). Moreover, more than half of all injured animals showed no allodynia at any level which ensures an important control group in studies of CNP to account for any physiological changes due to the injury itself.

Although a variety of methods have been used to identify CNP in rodent models of SCI, this study was primarily focused on identifying at-level allodynia evoked by a tactile stimulus. At-level allodynia is most often assessed in the rat by providing a sensory stimulus to the trunk and quantifying painful responses. For example, studies have examined the impact of a broad range of stimuli, including static mechanical (Baastrup et al., 2010; Crown et al., 2005, 2008; Hulsebosch et al., 2000; Lindsey et al., 2000), dynamic brushing or stroking, and/or gentle squeezing (Baastrup et al., 2010; Hall et al., 2010; Hubscher & Johnson, 1999). Although, a tactile stimulus is most often used to identify at-level allodynia, not all CNP must be evoked.

It is possible that animals experienced types of pain that cannot be evoked by a tactile stimulus, including pain evoked by other stimulus modalities (Finnerup et al., 2014; Siddall et al., 2003). Previous research on CNP has found the presence of cold (Lindsey et al., 2000; Yoon et al., 2004) and thermal hyperalgesia (Carlton et al., 2009; Putatunda et al., 2014). Therefore, now that a method has been developed to separate animals that develop tactile allodynia from those that do not, other stimulus modalities should be tested to obtain a more comprehensive understanding of the pain phenotypes that develop following a mid-thoracic SCI (Deuis et al., 2017).

In addition to stimulus evoked pain, it is possible that animals experienced spontaneous pain. Spontaneous pain has been assessed in rodents through the use of operant learning tasks like the place escape avoidance paradigm (PEAP), conditioned place preference, and mechanical conflict avoidance paradigm (Chhaya et al., 2019; Harte et al., 2016; Labuda & Fuchs, 2000; Sufka, 1994; Yang et al., 2014). However, these measures introduce confounding factors (e.g., motivation) that have yet to be resolved and may therefore interfere with pain assessment.

To refine the assessment of at-level allodynia, pain was precisely localized on the dorsum of the trunk. At-level allodynia was localized predominately at and just rostral to the site of the lesion (within dermatomes T4-T11), which is consistent with the girdle region previously reported (Baastrup et al., 2010; Hulsebosch et al., 2000; Lindsey et al., 2000). Interestingly, at-level allodynia in humans has been clinically defined as pain spanning up to three dermatome levels below and one dermatome above the level of the SCI, while below-level allodynia presents greater than three dermatomes below the level of the injury (Bryce et al., 2012). The relevance of this difference in the location of at-level allodynia between the rat model and the human is unclear, but could be due to differences in overall body size or the bipedal stance of humans compared to the quadrupedal stance of the rodent.

The precise localization of trunk allodynia allowed us to define a smaller grid and test a larger sample of animals. With this larger cohort, it was evident that at-level allodynia could be confidently detected by tracking only evoked vocalizations and avoidance behavior, consistent with earlier studies (Baastrup et al., 2010; Christensen & Hulsebosch, 1997; Hulsebosch et al., 2000; Lindsey et al., 2000; M’Dahoma et al., 2014). While it is also possible that animals experiencing pain vocalized at frequencies that are inaudible to the experimenters (Knutson et al., 2002), they have not been shown to be a superior indicator of pain than vocalizations that are emitted in the audible range (Williams et al., 2008). Therefore, solely monitoring vocalization and avoids should be sufficient to identify at level allodynia.

While the general location of at-level allodynia has been consistently reported without the use of a grid system, the use of the grid for pain testing on the trunk adds several benefits. Behaviorally, it allows consistency both within animals and across animals, important for studies which look at pain development over time. Additionally, the large grid can be used to adapt this method of pain testing to contusion injuries at different spinal levels, other pain modalities (heat, cold, etc.), or other pain models, allowing for comparisons across models. More importantly, the use of the grid system can be used to study the relationship between allodynia and hyperexcitability along the entire neural axis relative to the known somatotopy (Gwak & Hulsebosch, 2011). Within regions with specific somatotopy, knowing the dermatomes that are affected can improve the accuracy with which changes in excitability are measured, thereby providing greater insight into the mechanisms underlying the development of tactile allodynia. Thus, we conclude that a standardized grid should be used to uniformly assess the trunk region for allodynia.

For assessment of pain in the paws, our results further support the necessity of the detection of supraspinal responses. We found that over one-quarter of animals that developed hyperreflexia did not exhibit any supraspinal responses to paw stimulation. Consistent with Van Gorp et al. (2014), these results suggest that not all animals that develop hyperreflexia develop pain, therefore, the prevalence of pain is likely overestimated in studies that define pain based on a withdrawal threshold criterion alone. In this model of SCI, pain should be defined by the presence of supraspinal response because withdrawal of the paw is a spinal reflex, which may be altered due to injury, whereas supraspinal responses are brainstem reflexes which originate below the level of injury (Baastrup et al., 2010). Misclassifying animals into an experimental group considered to have pain could have substantial effects on the interpretation of results and the study of CNP in general.

Importantly, although allodynia may develop at any level, it is most likely to develop at-level in the mid-thoracic contusion model which makes this a good model to study the mechanisms underlying at-level allodynia. Notably, not every animal developed at-level allodynia. An advantage of this is that it allows for the study of pain mechanisms separate from injury. Moreover, it may be an advantage that the prevalence of above-level allodynia in this model was low, as it has been suggested that above level injury may not be due to the SCI itself (Cruz-Almeida et al., 2009), but rather it is associated with peripheral sensitization in response to secondary injury after SCI (Hulsebosch et al., 2009; Widerström-Noga, 2017). For these reasons, we suggest that the mid-thoracic contusion model is well-suited for the study of at-level pain.

Although the mid-thoracic contusion model is primarily utilized for the study of at-level allodynia, we report that trunk stimulation can also evoke spasticity of the hindlimbs without supraspinal responses. An SCI model that produces both allodynia and spasticity is clinically relevant because there is an increased prevalence of spasticity in SCI patients who experience CNP (Andresen et al., 2016). Moreover, in both allodynia and spasticity, although the evoking stimulus is tactile and not nociceptive, the stimulus locations that are evoked are not co-localized. Therefore, it is likely that allodynia and spasticity have different underlying mechanisms, but additional studies would be needed to confirm this.

Finally, the timing of the onset of allodynia was different depending on the pain phenotype. Consistent with human studies (Burke et al., 2017; Finnerup et al., 2014), we found that at-level allodynia had an earlier onset than below-level allodynia, suggesting that different mechanisms may be involved in development of allodynia in these different regions. Using this midthoracic contusion model and definition of at level allodynia, additional studies can probe potential differences by comparing animals with SCI that develop allodynia to those that do not.

In this study, we demonstrated that at-level tactile allodynia could be assessed in the rat mid-thoracic contusion model by quantifying the percent of stimulations that elicited vocalization and avoiding responses in a region of the trunk aligned to dermatomes T4-T11. This model produces a similar distribution of pain phenotypes observed in human SCI patients with the possibility of developing spasticity in response to tactile stimulation of thoracic dermatomes. This is important because associating unique pain phenotypes with underlying pathophysiology will be essential to the development of effective treatments for CNP.

## Acknowledgements

This work was supported by grant R01NS096971 to KAM.

## Author Contributions

G.H.B., B.N., A.K.S., and K.A.M. wrote the manuscript, G.H.B., B.N., A.K.S., and J.R. performed all experimental work, G.H.B., B.N., A.K.S. analyzed the data, G.H.B, B.N., M.R.D, J.R.B., and K.A.M. were involved in experimental design. All authors contributed to discussion and review of the results and the manuscript.

